# Integrating Predicted Transcriptome From Multiple Tissues Improves Association Detection

**DOI:** 10.1101/292649

**Authors:** Alvaro N. Barbeira, Milton D. Pividori, Jiamao Zheng, Heather E. Wheeler, Dan L. Nicolae, Hae Kyung Im

**Affiliations:** Section of Genetic Medicine, The University of Chicago, Chicago, IL, USA; Department of Biology, Loyola University Chicago, Chicago, IL, USA; Department of Computer Science, Loyola University Chicago, Chicago, IL, USA; Department of Statistics, The University of Chicago, Chicago, IL, USA; Department of Human Genetics, The University of Chicago, Chicago, IL, USA

## Abstract

Integration of genome-wide association studies (GWAS) and expression quantitative trait loci (eQTL) studies is needed to improve our understanding of the biological mechanisms underlying GWAS hits, and our ability to identify therapeutic targets. Gene-level association test methods such as PrediXcan can prioritize candidate targets. However, limited eQTL sample sizes and absence of relevant developmental and disease context restricts our ability to detect associations. Here we propose an efficient statistical method that leverages the substantial sharing of eQTLs across tissues and contexts to improve our ability to identify potential target genes: MulTiXcan. MulTiXcan integrates evidence across multiple panels while taking into account their correlation. We apply our method to a broad set of complex traits available from the UK Biobank and show that we can detect a larger set of significantly associated genes than using each panel separately. To improve applicability, we developed an extension to work on summary statistics: S-MulTiXcan, which we show yields highly concordant results with the individual level version. Results from our analysis as well as software and necessary resources to apply our method are publicly available.

**Author summary:** We develop a new method, MulTiXcan, to test the effect of gene expression regulation on complex traits, integrating information available across multiple tissue studies. We show this approach has higher power than traditional single-tissue methods. We extend this method to use only summary-statistics from public GWAS. We apply these methods to over 200 complex traits available in the UK Biobank cohort, and 100 complex traits from public GWAS and discuss the findings.

## Introduction

Recent technological advances allow interrogation of the genome to a high level of coverage and precision, enabling experimental studies that query the effect of genotype on both complex and molecular traits. Among these, GWAS have successfully associated genetic loci to human complex traits. GWAS meta-analyses with ever increasing sample sizes allow the detection of associated variants with smaller effect sizes [1–3]. However, understanding the mechanism underlying these associations remains a challenging problem, requiring follow-up studies and a wide array of techniques such as prioritization [4] and pathway analysis [5].

Another approach is the study of quantitative trait loci (eQTLs), measuring association between genotype and gene expression. These studies provide a wealth of biological information but tend to have smaller sample sizes. A similar observation applies to QTL studies of other traits such methylation, metabolites, or protein levels.

The importance of gene expression regulation in complex traits [6–9] has motivated the integration of eQTL studies and GWAS. To examine these mechanisms we developed PrediXcan [10], a method that tests the mediating role of gene expression variation in complex traits. We also developed an extension that accurately infers its gene-level association results using summary statistics data: S-PrediXcan [11]. This allows the rapid exploration of information available in publicly available GWAS summary statistics results, at a dramatically reduced computational burden.

Due to sharing of eQTLs across multiple tissues, we have shown the benefits of an agnostic scanning across all available tissues [11]. Despite the increased multiple testing burden (for Bonferroni correction, the total number of gene-tissue pairs must be used when determining the threshold) we gain considerably in number of significant genes. However, given the substantial correlation between different tissues [12], Bonferroni correction can be too stringent increasing the false negative rate.

In order to aggregate evidence more efficiently, here we present a method termed MultiXcan that tests the joint effects of gene expression variation from different tissues. Furthermore, we develop and implement a method that only needs summary statistics from a GWAS: Summary-MulTiXcan (S-MulTiXcan for short). We make our implementation publicly available to the research community in https://github.com/hakyimlab/MetaXcan. We apply this method to traits from the UK Biobank study and over a hundred public GWAS, and publish the results in http://gene2pheno.org.

## Results

### Combining Information Across Tissues Through Multivariate Regression

To combine information across tissues, we regress the phenotype of interest on the predicted expression of the gene in multiple tissues as follows:

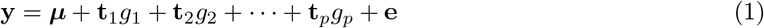

where y is the *n*-dimensional phenotype vector, ***μ*** is an intercept term, **t**_*i*_ is predicted expression of the gene in tissue *i, g_i_* is its effect size, and **e** an error term with variance 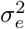; *p* is the number of available tissue models.

Expression predictions from many of these tissues are highly correlated. To avoid numerical issues caused by collinearity, we compute the principal components of the predicted expression matrix and discard the axes of smallest variation. Additional covariates can be added to the regression seamlessly. Fig 1-a displays an overview of the method; see further details in the Methods section. We illustrate prediction correlation across models in Supp. Fig 1.

**Figure 1.**
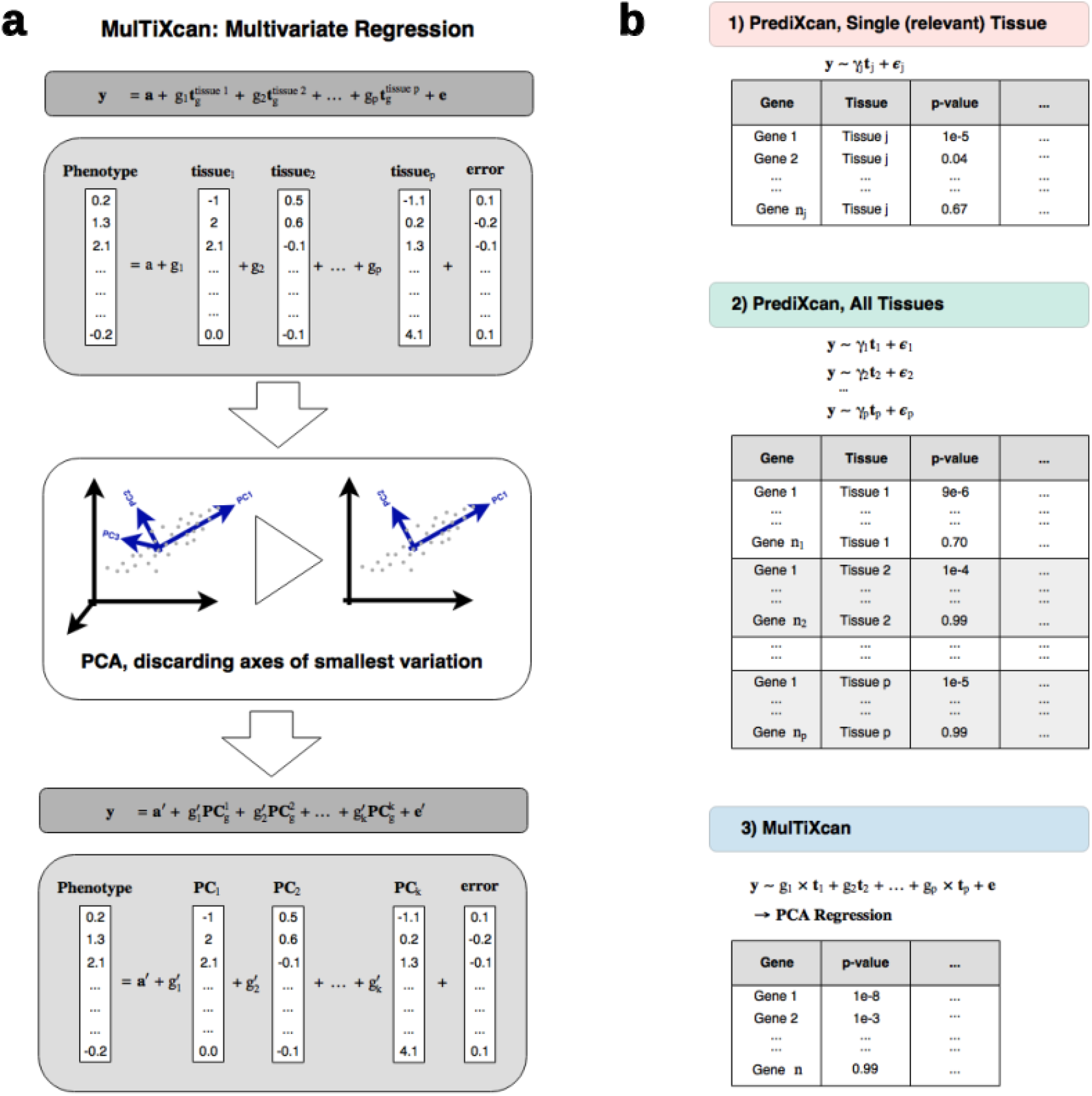
MulTiXcan method. **Panel a** illustrates MulTiXcan method. Predicted expression from all available tissue models are used as explanatory variables. To avoid multicolinearity, we use the first k Principal Components of the predicted expression. y is a vector of phenotypes for *n* individuals, 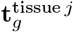 is the standardized predicted gene expression for tissue *j*, g_*j*_ is its effect size, a is an intercept and e is an error term. **Panel b** shows a schematic representation of MulTiXcan results compared to classical PrediXcan, both for a single relevant tissue and all available tissues in agnostic scanning. y is a vector of phenotypes for *n* individuals, **t**_*j*_ is the standardized predicted gene expression for model *j*, *g_j_* is its effect size in the joint regression, *γ_j_* is its effect size in the marginal regression using only prediction *j*, e and *є_j_* are error terms.

### MulTiXcan Detects More Associations Than Single-Tissue PrediXcan

We applied our method to 222 traits from UK Biobank [13]. We used prediction models for 44 tissues trained with Genotype-Tissue Expression (GTEx) samples [12]. We found that it can detect many more associations than PrediXcan using a specific tissue or even when aggregating results from all tissues (Bonferroni-corrected for all gene-tissue pairs tested).

More specifically, we compared three approaches for assessing the significance of a gene jointly across all tissues: 1) running PrediXcan using the most relevant tissue; 2) running PrediXcan using all tissues, one tissue at a time; 3) running MulTiXcan. Fig 1-b illustrates a schematic representation of the results from each approach. Overall, MulTiXcan detects more associations than PrediXcan, as shown in Fig 2-c. Supplementary Data 1 contains a summary of associations per trait. See Supplementary Data 2 and 3 for the list of significant MulTiXcan and PrediXcan results respectively.

**Figure 2.**
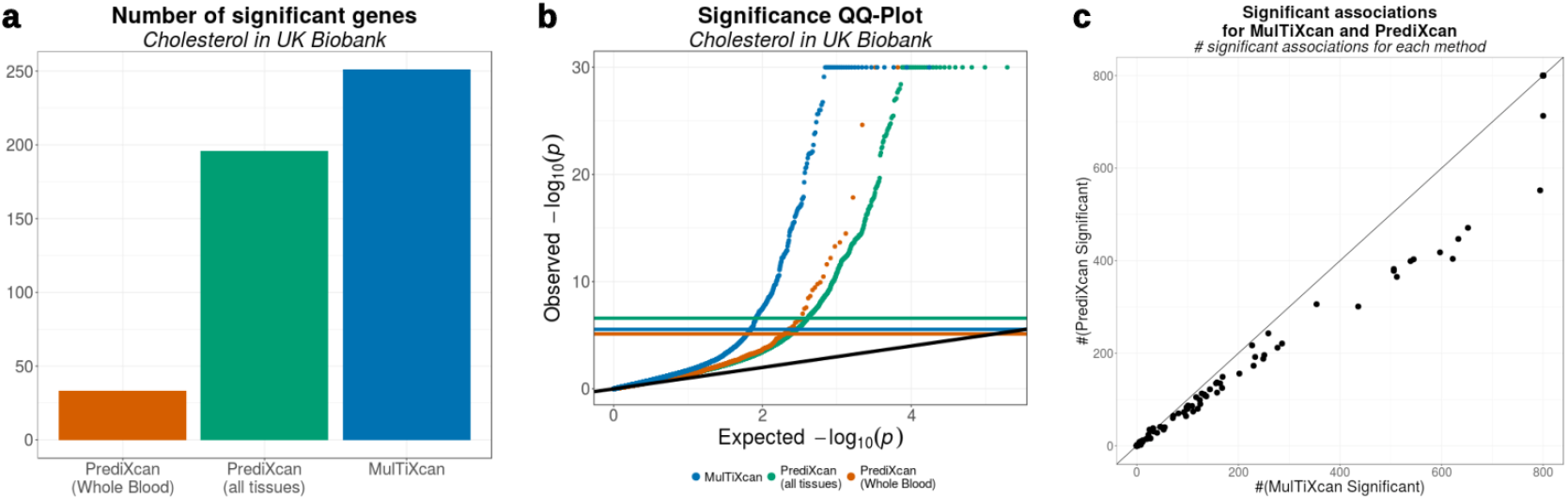
Joint testing across all tissues increases number of significant genes. **Panel a** shows the number of discoveries in each method for Cholesterol trait. MulTiXcan is able to detect more findings (251 significant associations) than either of PrediXcan approaches (33 using only Whole Blood and 196 using all 44 GTEx tissues). **Panel b** compares the distribution of MulTiXcan’s p-values to PrediXcan’s p-values for the Cholesterol trait in the UK Biobank cohort. Both PrediXcan with a single tissue model (GTEx Whole Blood) and 44 models (GTEx v6p models) are shown. Notice that Bonferroni-significance levels are different for each case, since 6588 genes were tested in PrediXcan for Whole Blood, 195532 gene-tissue pairs for all GTEx tissues, and 17434 genes in MulTiXcan. P-values were truncated at 10^-30^ for visualization convenience. **Panel c** compares the number of significant associations discovered by MulTiXcan and PrediXcan for 222 traits from UK Biobank. These numbers were thresholded at 800 for visualization purposes.

We examined the High-Cholesterol trait results in closer detail. We used 50,497 cases and 100,994 controls. After Bonferroni correction, MulTiXcan was able to detect a larger number of significantly associated genes (251) than PrediXcan using all tissues (196) or only a single tissue (whole blood, 33). 172 genes were detected by both PrediXcan and MulTiXcan. Fig 2-a compares the number of significantly associated genes in MulTiXcan, and PrediXcan both for a single tissue (whole blood) and all tissues. Fig 2-b shows the QQ-plot for associations in these three approaches. There are 79 genes associated to high cholesterol via MulTiXcan and not PrediXcan. Among them, we find many genes related to lipid metabolism (*APOM* [14], *PAFAH1B2* [15]), glucose transport(*SLC5A6* [16]), and vascular processes (*NOTCH4* [17]).

### MulTiXcan Results Can Be Inferred From GWAS Summary Results

To expand the applicability of our method to massive sample sizes and to studies where individual level data are not available, we have extended our method to use GWAS summary results rather than individual-level data.

We call this extension Summary-MulTiXcan (S-MulTiXcan). Here we derive an analytic expression that relates the joint regression estimates (*g_j_*) to the marginal regression estimates (*γ_j_* as obtained from S-PrediXcan), assuming a known LD structure from a reference panel. We display a conceptual overview of the method in Fig 4-a. See details in the Methods Section.

Fig 3 displays a few examples of the general agreement between the individual-level MulTiXcan and S-MulTiXcan. The summary-based version’s results tend to be slightly less significant than MulTiXcan, as illustrated in Supp. Fig 2.

**Figure 3.**
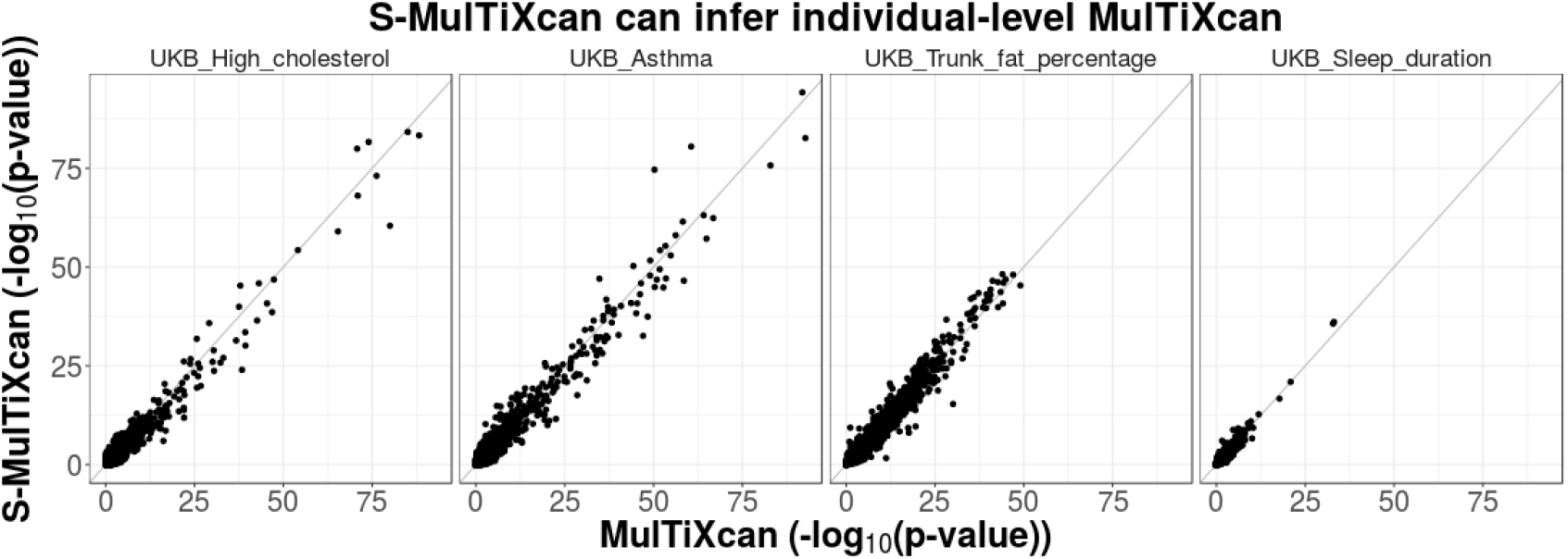
MulTiXcan results can be inferred from GWAS summary statistics and a reference panel. This figure compares S-MulTiXcan to MulTiXcan in three UK Biobank phenotypes. GTEx individuals were used as a reference panel for estimating expression correlation in the study population. The summary data-based method shows a good level of agreement with the individual-based method. In cases where the LD-structure between reference and study cohorts is mismatched, the summary-based method becomes less accurate. For example in Asthma, two genes are significantly overestimated; however it tends to be conservative for most genes.

To reduce false positives due to LD misspecification, we discard any significant association result for a gene when the best single tissue result has p-value greater than 10^−4^. This is rather conservative since it is possible that evidence with modest significance from weakly correlated tissues can lead to very significant combined association. We have, in fact, found several such instances using individual level data.

### Application to a broad set of complex traits

We applied S-MulTiXcan to over a 100 traits on publicly available GWAS. As with the individual level method, we observed that S-MulTiXcan detects more associations than S-PrediXcan in most cases, as shown in Fig 4-b, after discarding suspicious associations. We also show the QQ-plots and total number of detected association for a sample trait (Schizophrenia) on Fig 4-c and 4-d.

**Figure 4.**
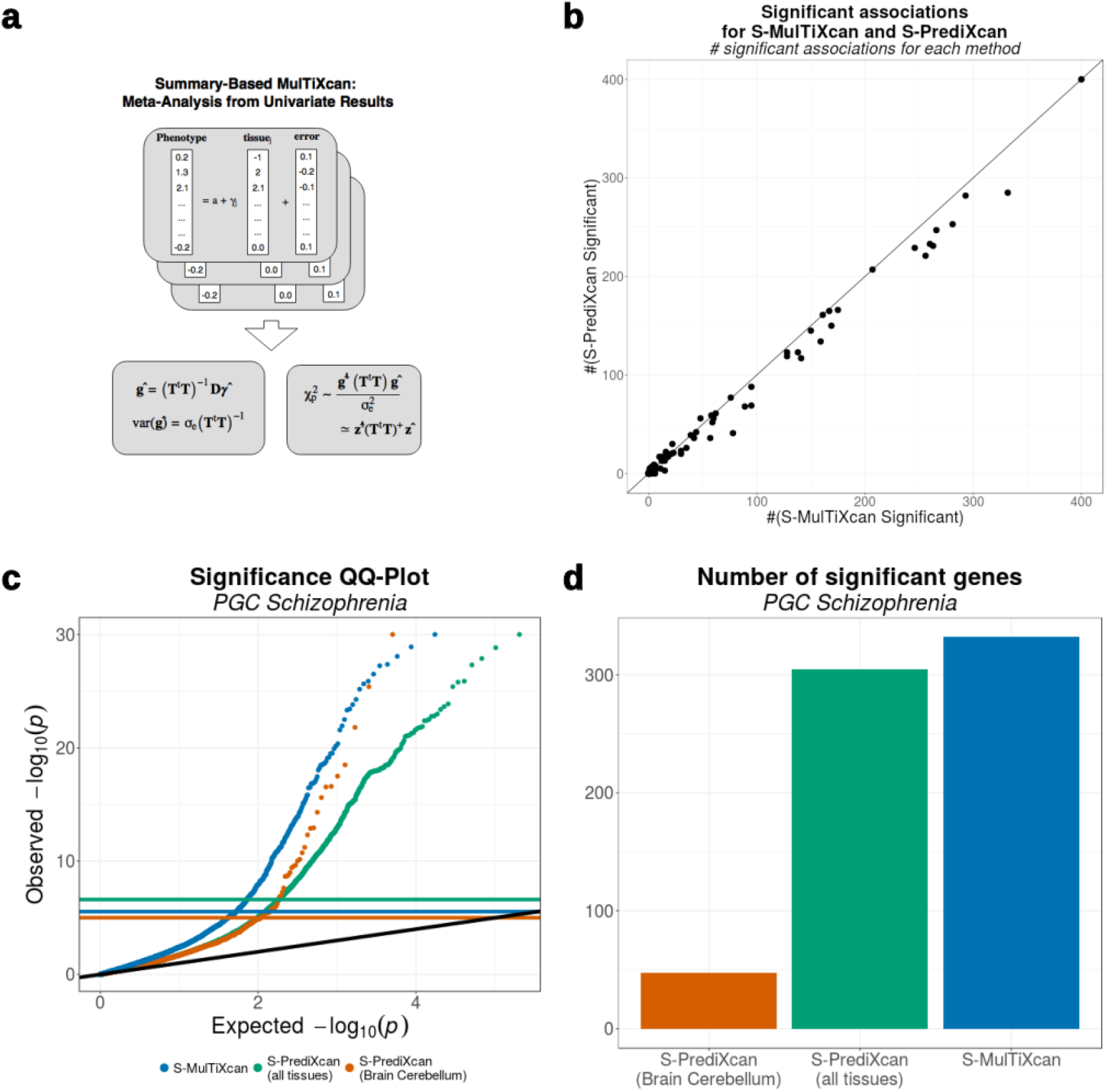
Comparison between S-PrediXcan and S-MulTiXcan. **Panel a** illustrates the S-MulTiXcan method: the marginal univariate S-PrediXcan effect sizes are computed, then the joint effect sizes are estimated from them. The significance is quantified through an omnibus test. **Panel b** compares the number of associations significant via S-MulTiXcan versus those significant via S-PrediXcan, for the same GWAS Studies. In most cases, S-MulTiXcan detects a larger number of exclusive significant associations. The number of discoveries was thresholded at 200 for visualization purposes. **Panel c** displays QQ-Plots for the association p-values from S-MulTiXcan and S-PrediXcan in Schizophrenia, using a model trained on brain’s cerebellum, and S-PrediXcan associations for all 44 GTEx tissues. **Panel d** shows the number of significant associations in Schizophrenia for each method as a bar plot.

These results have been incorporated into the publicly available catalog at http://gene2pheno.org. The list of analyzed traits can be found in Supplementary Data 4. Supplementary Data 5 contains a summary of significant associations for each trait. Supplementary Data 6 lists the significant S-MulTiXcan results for each trait.

### Highlight Of Associations Identified By Summary-MulTiXcan

We examined the biological relevance of some of the genes detected by our new method that was missed by looking at one tissue at a time (S-PrediXcan).

For example, in the Early Growth Genetics (EGG) Consortium’s Body-Mass Index (BMI) study, S-MulTiXcan detects three genes not significant in S-PrediXcan: *POMC* (p-value=1.4 × 10^−6^, tied to childhood obesity [18]); *RACGAP1* (p-value= 1.2 × 10^−10^; embryogenesis [19], cell growth and differentiation, [20]); and *TUBA1B* (p-value= 1.23× 10^−09^, circadian cycle processes and psychological disorders [21], suggesting a behavioral pathway).

In the CARDIoGRAM+C4D Coronary Artery Disease (CAD) study, S-MulTiXcan detected 12 associations not significant in S-PrediXcan. The top result was *AS3MT* (p-value=4.3 × 10^−9^), related to arsenic metabolism; interestingly, environmental and toxicological studies link arsenic exposure and *AS3MT* polymorphisms with cardiovascular disease [22,23]. Associations previously linked to CAD included *CDKN2B* (p-value< 1.0 × 10^−6^, [24]) *HECTD4* (p-value< 2.3 × 10^−6^, [25]). Other interesting S-MulTiXcan findings were *CLCC1* (pvalue=1.2 × 10^−7^, a gene for chloride channel activity); *IREB2* (p-value=2.1 × 10^−7^, recently linked to pulmonary conditions, [26]), and *ADAM15* (p-value=2.5 × 10^−07^, from the disintegrin and metalloproteinase family, linked to atherosclerosis [27], atrial fibrillation [28], and other vascular processes [29,30]).

The list of significant S-MulTiXcan and S-PrediXcan results for all traits can be found in Supplementary Data 6 and 7.

## Discussion

Motivated by the widespread sharing of regulatory processes across tissues [12], we propose here a method (MulTiXcan) that aggregates information across multiple tissues and improves the identification of genes significantly associated with complex traits. To expand the applicability of our approach we have extended the method to accommodate GWAS studies where only summary results are available (S-MulTiXcan). We show through applications to hundreds of traits the performance of both individual and summary based methods. We also show that the summary based method provides a reasonably good approximation to the individual level results.

As any method relying on a reference panel, S-MulTiXcan might be inaccurate when the study population has a different Linkage Disequilibrium (LD) structure than the reference panel. For example, should two models for a gene yield predicted expressions that are lowly correlated in the reference panel but highly correlated in the study population, then this method underestimates their correlation. To avoid this misspecification, a reference panel matching the study population should be used when available (i.e. using East Asian population from 1000 Genomes if the study set is composed of East Asian individuals).

A limitation of PrediXcan and S-PrediXcan is LD contamination, i.e. when causal loci for the trait and expression are different but in LD. We have addressed this in S-PrediXcan through an additional colocalization filtering step. For MulTiXcan, this could be avoided by restricting the analysis to gene-tissue pairs with high colocalization probability.

Here we showed the advantages of our joint estimation method through application to multiple traits with publicly available GWAS results as well as the ones available in the UK Biobank. These results include many novel associations of interest, which we make publicly available to the research community in http://gene2pheno.org.

### Software And Resources

We make our software publicly available on a GitHub repository: https://github.com/hakyimlab/MetaXcan. Prediction model weights and covariances for different tissues can be downloaded from PredictDB.org. A short working example can be found on the GitHub page; more extensive documentation can be found on the project’s wiki. The results of S-MulTiXcan applied to the 44 human tissues and a broad set of phenotypes can be queried on http://gene2pheno.org.

## Methods

### Definitions, Notation And Preliminaries

Let us consider a GWAS study of *n* samples, and assume availability of prediction models in *p* different tissues. Each model *j* is a collection of prediction weights 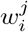.

Let:

- y be an *n*-vector of phenotypes, assumed to be centered for convenience.
- **X** the genotype matrix, where each column *X_l_* is the *n*-vector genotype for SNP *l*. We assume it coded in the range [0,2] but it can be defined in another range, or standardized.
- 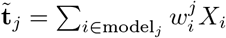 be the predicted expression in tissue *j*. This is the independent variable used by single-tissue PrediXcan.
- **t**_*j*_ be the standardization of 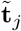 to *mean* = 0 and *standard deviation* = 1.

In our application, we will consider *p* = 44 models for a given gene’s expression, trained on GTEx data. This method is easily extensible to support incorporation of other covariates, or correction by them.

### MulTiXcan

MulTiXcan consists of fitting a linear regression of the phenotype on predicted expression from multiple tissue models jointly:

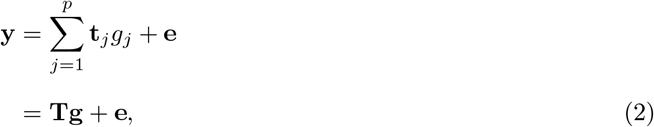

where y is a vector of phenotypes for *n* individuals, **t**_*j*_ is an *n*-vector of standardized predicted gene expression for model *j, g_j_* is the effect size for the predicted gene expression *j*, e is an error term with variance 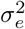 and *p* is the number of tissues; thus **T** is a data matrix where each column *j* contains the values from **t**_*j*_, and g is the *p*-vector of effect sizes *g_j_*. One of this columns is a constant intercept term.

The high degree of eQTL sharing between different tissues induces a high correlation between predicted expression levels. In order to avoid collinearity issues and numerical instability, we decompose the predicted expression matrix into principal components and keep only the eigenvectors of non negligible variance. To select the number of components, we used a condition number threshold of 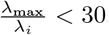, where λ_*i*_ is an eigenvalue of the matrix **T**^*t*^**T**. A range of values between 10 and 100 yielded similar results for significance in real data. See the next section for additional details in the number of components used. Lastly, we use an F-test to quantify the significance of the joint fit.

We use Bonferroni correction to determine the significance threshold. For MulTiXcan, we use the total number of genes with a prediction model in at least one tissue, which yields a threshold approximately at 0. 05/17500 ~ 2.9 × 10^−6^. For PrediXcan across all tissues, we use the total number of gene-tissue pairs, which yields a threshold approximately at 0.05/200, 000 ~ 2.5 × 10^−7^.

### Application To UK Biobank Data

We used the same covariates reported in [31], which include the first ten genotype principal components, sex, age, genotyping array, and depending on the trait others such as body mass index (BMI), weight or height. We used 44 models trained on GTEx tissues from release version v6p. For diseases, we used twice as many healthy individuals as controls, selected at random. For the MulTiXcan-significant associations in the 222 traits, the median number of available models is 11 (1*Q* = 7, 3*Q* = 16), with ~ 77% components surviving PCA thresholding.

### Summary-MulTiXcan

We have demonstrated that S-PrediXcan can accurately infer PrediXcan results from GWAS Summary Statistics and LD information from a reference panel [11], with the added benefits of reduced computational and regulatory burden. Here we extend MulTiXcan in a similar fashion.

Summary-MulTiXcan (S-MulTiXcan) infers the individual-level MulTiXcan results, using univariate S-PrediXcan results and LD information from a reference panel. It consists of the following steps:

- Computation of single tissue association results with S-PrediXcan.
- Estimation of the correlation matrix of predicted gene expression for the models using the Linkage Disequilibrium (LD) information from a reference panel (typically GTEx or 1000 Genomes [32])
- Discarding components of smallest variation from this correlation matrix to avert collinearity and numerical problems (Singular Value Decomposition, analogue to PC analysis in individual-level data).
- Estimation of joint effects from the univariate (single-tissue) results and expression correlation.
- Discarding suspicious results, suspect to be false positives arising from LD-structure mismatch.

### Joint Analysis Estimation From Marginal Effects

To derive the multivariate regression (2) effect sizes and variances using the marginal regression (3) estimates, we employ a technique presented in [33].

More specifically, we want to obtain the multivariate regression coefficient estimates for *g_j_* (2) using the estimates from the marginal regression:

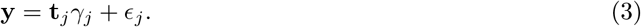

where we assume y centered for convenience, and *є_j_* is the marginal regression error term with variance 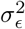 (i.e. we assume a common variance 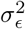 for all *j*).

First, notice that the solution to the multivariate regression in Eq (2) is

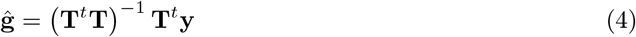

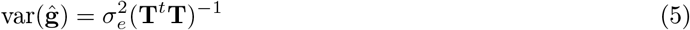

whereas the solution to the marginal regression in Eq (3) is:

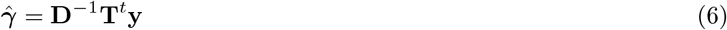

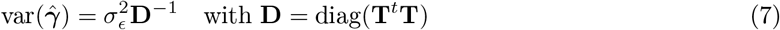

where *γ* is the vector of effect sizes *γ_j_*, and 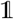 is the *p* × *p* identity matrix. Please note that, since the **t**_*j*_ are standardized, then 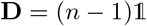 and 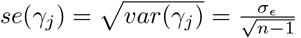.

From (6) we get 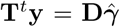, which we replace in (4) and obtain the relationship between marginal and joint estimates:

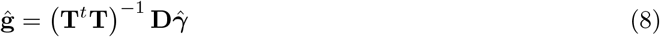

To compute the variance of the estimated effect sizes (5) we use the variance of the phenotype as a conservative estimate of 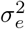 and LD information from reference samples as described next.

### Estimating Expression Correlation From A Reference Panel

As the genotypes from most GWAS are typically unavailable, we must use a reference panel to compute **T**^*t*^**T**, using only those SNPS available in the GWAS results. To do so, notice that:

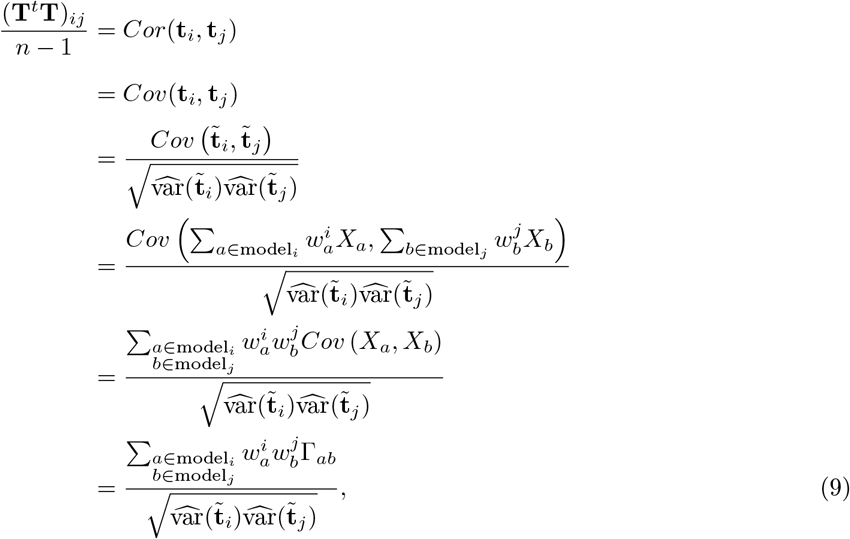

where Γ_*ij*_ are the elements of the covariance matrix 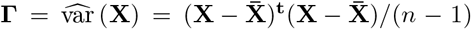. We compute the variances as in the S-PrediXcan analysis:

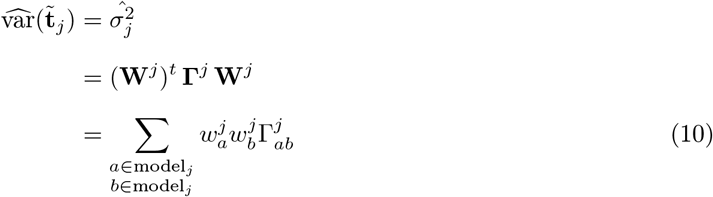

### Addressing Singularity Of The Correlation Matrix

Given the high degree of correlation among many of the prediction models, **T**^*t*^**T** is close to singular and its inverse cannot be reliably calculated for many genes. To address this problem, we compute the pseudo-inverse via Singular Value Decomposition, decomposing the correlation matrix into its principal components and removing those with small eigenvalues. In other terms, we will restrict the analysis to axes of largest variation of the expression data. This is analogous to the principal components-based approach used with individual level data. We denote with **Σ**^+^ the pseudo-inverse for any matrix **Σ**. We use the same condition number from individual-level MultiXcan 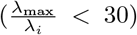 as threshold. For S-MulTiXcan-significant associations across 100 public traits, we found a median number of available models of 9 (1*Q* = 5, 3*Q* = 15), with ~ 80% of components surviving the SVD threshold.

### Estimating Significance

To quantify significance, we use the fact that the regression coefficient estimates follow a (approximate) multivariate normal distribution: 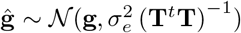. Under the null hypothesis of no association, it follows that 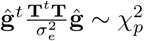. We can then replace g with its estimate from the marginal regression:

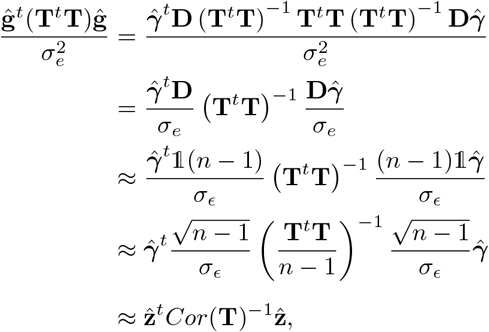

where *Cor*(**T**) is the autocorrelation of **T**, and 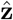 is the *p*-vector of marginal analysis z-scores, *γ_j_*/*se*(*γ_j_*). We have used 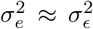 as an approximation (i.e. the residual variance of the *marginal* regression as approximation of the residual variance of the *joint* regression). This simplification is conservative, and based on our comparison to the individual multivariate results we consider the loss of efficiency acceptable.

In practice, we will use the SVD pseudo-inverse *Cor*(**T**)^+^ as explained in the previous section, and a 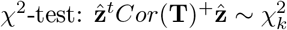, with *k* the number of components surviving the SVD pseudoinverse.

### Implementation And Computation

Prediction Models were obtained from PredictDB.org resource. These models were trained using Elastic Net as implemented in R’s package *glmnet* [34], with a mixing parameter *α* = 0.5, over 44 tissue studies from GTEx’ release version 6p. The underlying GTEx study data was obtained from dbGaP with accesion number phs000424.v6.p1. Please see [11] for details. We implemented MulTiXcan and S-MulTiXcan working up from existing software in the MetaXcan package.

UK Biobank genotype data for 487,409 individuals was downloaded and processed in the Bionimbus Protected Data Cloud (PDC), a secure biomedical cloud operated at FISMA moderate as IaaS with an NIH Trusted Partner status for analyzing and sharing protected datasets. We computed GWAS results using BGENIE, a program for efficient GWAS for multiple continuous traits [35]. We selected 222 traits available for these individuals, covering continuous phenotypes such as height and self reported diseases such as asthma. We used different covariate groups for these phenotypes as in [31]. Age, sex and the top ten principal components were used in all cases. For diseases, we randomly sampled twice as many healthy controls as there were cases. Gene expression prediction was computed on the genotype data using the 44 GTEx models.

When running MulTiXcan, we used the same covariates and data as in the GWAS. On most continuous phenotypes, there were between 300,000 and 400,000 individuals with available data determined by the intersection of covariates and traits. For the case of self reported diseases, we found a number of cases ranging from a few hundreds (i.e. Acne) to 50,000 (i.e. High Cholesterol). We also ran S-PrediXcan on 105 public GWAS traits (the same analyzed in [11], see Supplementary Data 4 for details).

## Supplementary Data

### Supplementary Data 1. Summary statistics for UK Biobank traits used in the MulTiXcan analysis

MulTiXcan was run for 222 traits on UK Biobank. Summary statistics for significant results included in **supp-data-ukb-multixcan-stats.txt**. Columns are: **tag**: trait, gene2pheno.org display name; **n_predixcan_significant**: Number of Bonferroni-significant PrediXcan results; **n_MulTiXcan_significant** number of Bonferroni-significant results for MulTiXcan; **n_predixcan_only** number of results only significant in PrediXcan; **n_MulTiXcan_only** number of results only significant in MulTiXcan.

### Supplementary Data 2. Significant associations for MulTiXcan on UK Biobank

Significant results included in **supp-data-ukb-multixcan-significant.txt**. Columns are: **phenotype**: trait, gene2pheno.org display name; **gene**: Ensembl id; **gene_name**: HUGO name; **pvalue**: p-value of the S-MulTiXcan association; **n_models** number of prediction models available for the gene; **n_used** number of independent components surviving PCA selection; **n_samples**: number of individuals available.

### Supplementary Data 3. Significant associations for PrediXcan on UK Biobank

Significant results included in **supp-data-ukb-p-significant.txt**. Columns are: **Phenotype**: trait, gene2pheno.org display name; **model**: GTEx tissue where the model was trained; **gene**: Ensembl Id; **gene_name**: HUGO name; **model** GTEx tissue where model was trained; **zscore** PrediXcan association Z-score, **pvalue** PrediXcan association p-value; **n_samples**: number of individuals available.

### Supplementary Data 4. List of Genome-wide Association Meta Analysis (GWAMA) Consortia and phenotypes

Data included in **supp-data-gwas-traits.txt**. Columns are consortium name, study name, gene2pheno.org display name, study sample size, study population, URL of portal where data was downloaded from, link to pubmed entry if available.

### Supplementary Data 5. Summary statistics for traits used in the MulTiXcan analysis

MulTiXcan was run for 105 public GWAS. Summary statistics for significant results included in **supp-data-gwas-smultixcan-stats.txt**. Columns are: **tag**: gene2pheno.org display name; **consortium**: Consortium Name; **name**: study name; **n_spredixcan_significant**: Number of Bonferroni-significant S-PrediXcan results; n_sMulTiXcan_significant number of Bonferroni-significant results for MulTiXcan; **n_spredixcan_only** number of results only significant in S-PrediXcan; **n_sMulTiXcan_only** number of results only significant in S-MulTiXcan.

### Supplementary Data 6. Significant associations for Summary-MulTiXcan on public GWAS

Significant results included in **supp-data-gwas-smultixcan-significant.txt**. Columns are: **tag**: gene2pheno.org display name; consortium: **Consortium** Name; **name**: study name; **gene**: Ensembl id; **gene_name**: HUGO name; **pvalue**: p-value of the S-MulTiXcan association; **n** number of S-PrediXcan results available for the gene; **n_indep** number of independent components surviving SVD; **p_i_best** best p-value of S-PrediXcan;**t_i_best** tissue that presented best S-PrediXcan result**; p_i_worst** worst p-value of S-PrediXcan; **t_i_worst** tissue that presented worst S-PrediXcan result; **suspicious**: whether the result was discarded as a potential false positive.

### Supplementary Data 7. Significant associations for Summary-PrediXcan on public GWAS

Significant results included in **supp-data-gwas-sp-significant.txt**. Columns are: **consortium**: Consortium Name; **name**: study name; **tag**: gene2pheno.org display name; **gene**: Ensembl Id; **gene_name**: HUGO name; **model** GTEx tissue where model was trained; **zscore** S-PrediXcan association Z-score, **pvalue** S-PrediXcan association p-value.

---

**Supplementary Figure 1.**
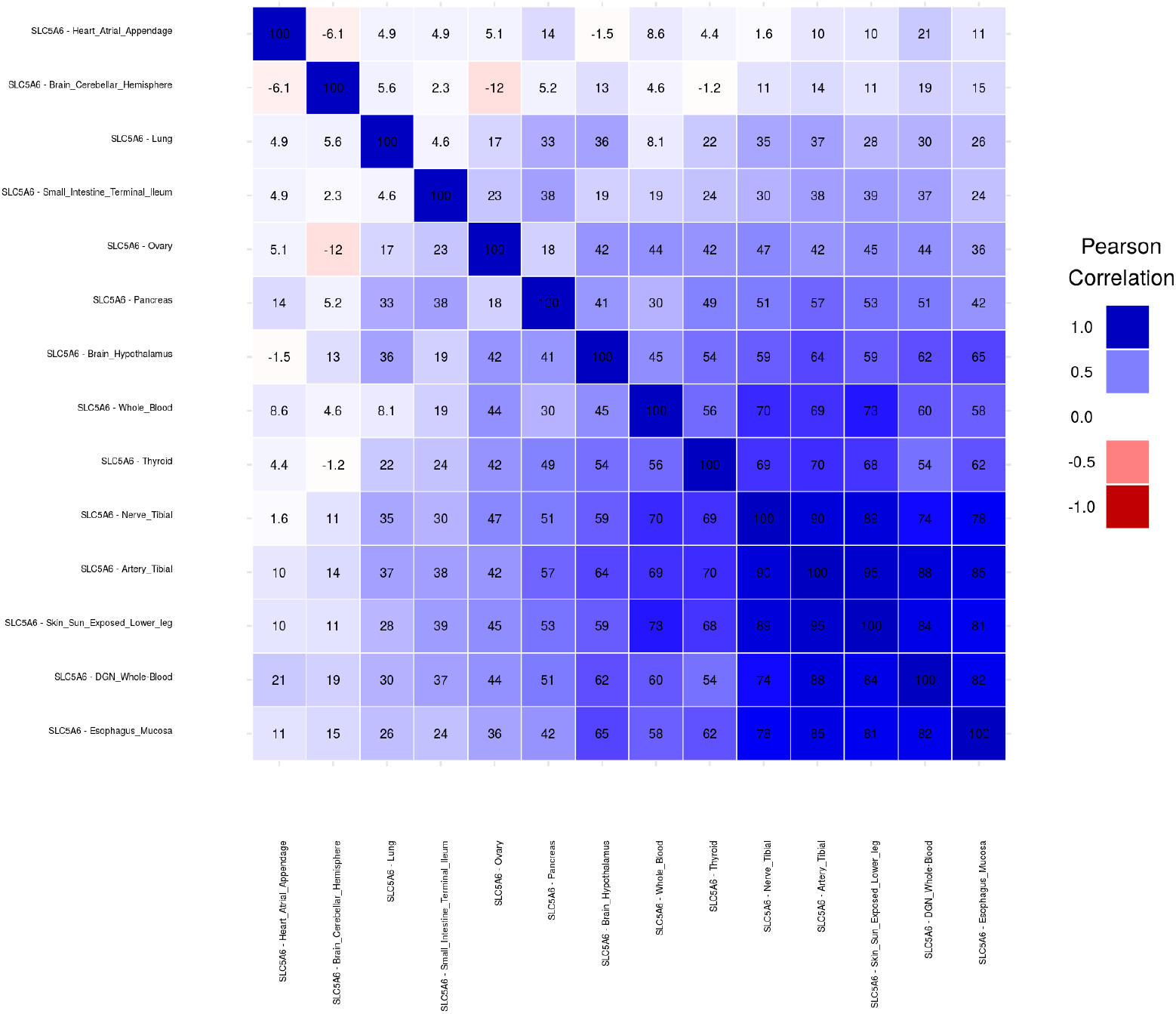
Predicted expression correlation for gene *SLC5A6*. We observe a high degree of predicted expression correlation, in agreement with recent publications on the high degree of mechanism sharing across tissues [12]. This behavior is exhibited in most genes.

**Supplementary Figure 2.**
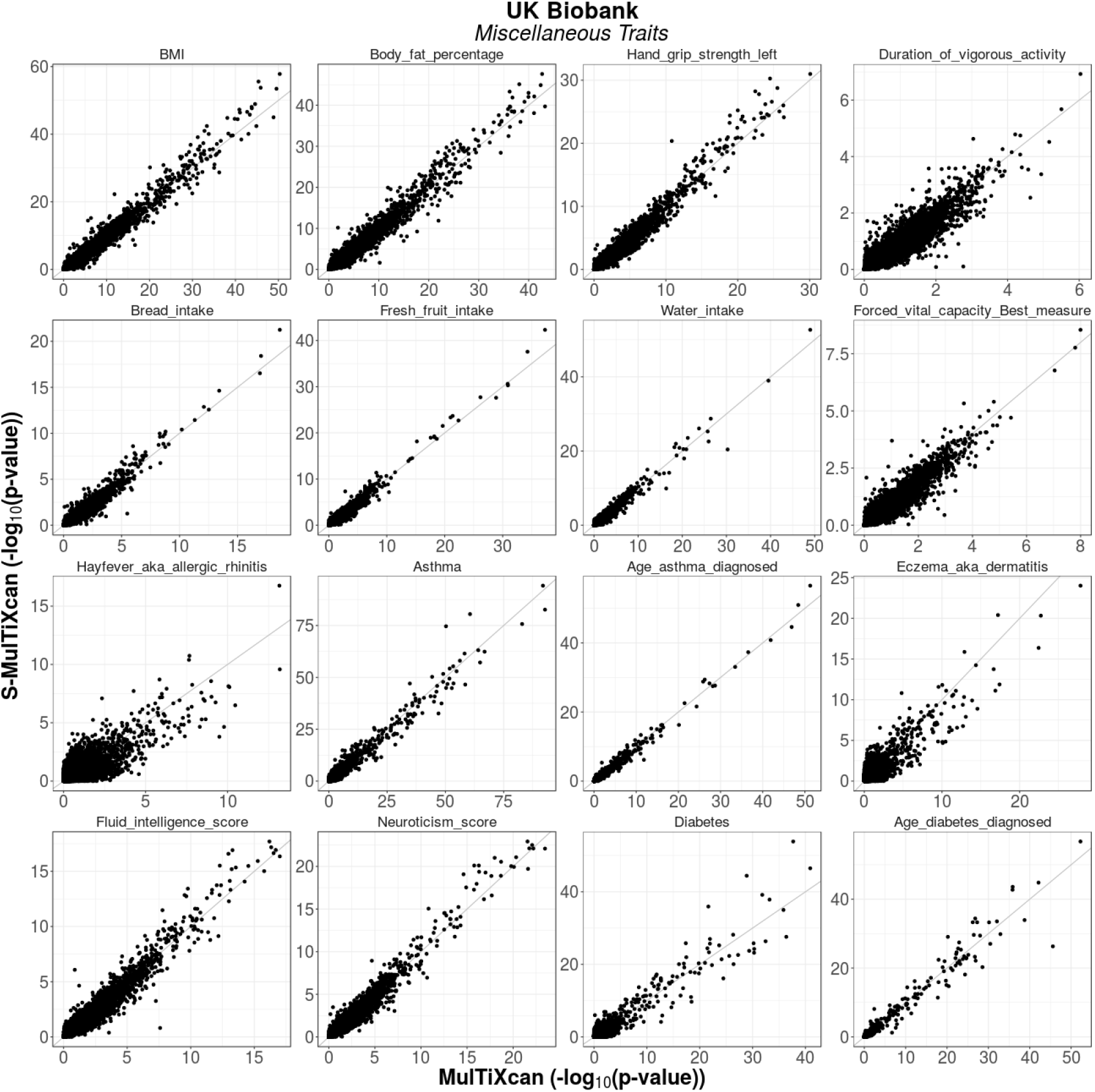
Summary-MulTiXcan vs MulTiXcan for Miscellaneous Traits. There is a satisfactory agreement between the individual-level and the summary-level versions of MulTiXcan in UK Biobank traits.

